# ProT-Diff: A Modularized and Efficient Approach to De Novo Generation of Antimicrobial Peptide Sequences through Integration of Protein Language Model and Diffusion Model

**DOI:** 10.1101/2024.02.22.581480

**Authors:** Xue-Fei Wang, Jing-Ya Tang, Han Liang, Jing Sun, Sonam Dorje, Bo Peng, Xu-Wo Ji, Zhe Li, Xian-En Zhang, Dian-Bing Wang

## Abstract

Antimicrobial Peptides (AMPs) represent a promising class of antimicrobial agents crucial for combating antibiotic-resistant pathogens. Despite the emergence of deep learning approaches for AMP discovery, there remains a gap in efficiently generating novel AMPs across various amino acid lengths without prior knowledge of peptide structures or sequence alignments. Here we introduce ProT-Diff, a modularized and efficient deep generative approach that ingeniously combines a pre-trained protein language model with a diffusion model to de novo generate candidate AMP sequences. ProT-Diff enabled the rapid generation of thousands of AMPs with diverse lengths within hours. Following in silico screening based on physicochemical properties and predicted antimicrobial activities, we selected 35 peptides for experimental validation. Remarkably, 34 of these peptides demonstrated antimicrobial activity against Gram-positive or Gram-negative bacteria, with 6 exhibiting broad-spectrum efficacy. Of particular interest, AMP_2, one of the broad-spectrum peptides, displayed potent antimicrobial activity, low hemolysis, and minimal cytotoxicity. Further in vivo assessment revealed its high effectiveness against a clinically relevant drug-resistant *E. coli* strain in a mouse model of acute peritonitis. This study not only presents a viable generative strategy for novel AMP design but also underscores its potential for generating other functional peptides, thereby broadening the horizon for new drug development.

## Introduction

The emergence of antibiotic-resistant pathogens has become one paramount public health concern, posing a significant threat to human well-being. However, since the 1990s, the pace of development and commercialization of new antibiotics have notably slowed down, with only a limited number of antimicrobial therapies receiving approval from regulatory agencies like the U.S. Food and Drug Administration (FDA) and the European Medicines Agency (EMA)^1^. Therefore, there is an urgent need to develop efficient approaches to identify alternative molecules to combat drug resistance.

Antimicrobial peptides (AMPs), a diverse group of peptides found across all life forms, play a vital role in the innate immune response by eradicating various microbes, including antibiotic-resistant micro-organisms^2, 3^. These peptides eliminate pathogens through diverse mechanisms, including mediating membrane disruption, causing DNA degradation, interfering with oxygen uptake ^4–6^. Furthermore, because their diminished risk of fostering antimicrobial resistance, AMPs have garnered significant attention as next-generation antibiotics^7–9^. Over the past few decades, numerous AMPs have been identified from natural sources, primarily through experimental methods, with some progressing to clinical trials^10^. However, limitations persist regarding their efficacy, stability, and toxicity, hindering their broad clinical applicability. Consequently, several advanced high-throughput techniques have been developed to discover new AMPs with enhanced performance^11, 12^. However, these experiment-driven approaches are time-consuming and costly, significantly limiting the efficiency of searching for desired drug candidates.

Recent advances in artificial intelligence (AI)-based approaches have greatly facilitated the discovery of AMPs. For instance, a predictive pipeline that integrates multiple natural language processing neural network models has been developed to discriminate AMPs from human gut microbiome^13^. This approach efficiently explores vast sequence datasets with reduced time required for mining novel functional peptides, but still faces challenges in generating peptide sequences that do not exist in nature. To address this limitation, a unified pipeline identified three unnatural hexapeptides with excellent antibacterial activity by extensively exploring the sequence space within a pre-constructed dataset of 6–9 amino acids^14^. However, this pipeline has to use a pre-constructed database, which may significantly limit the explored search space, thereby restricting the length of generated AMPs. Moreover, deep generative neural networks, such as Variational Autoencoders (VAE), Long Short-Term Memory (LSTM) and their variants, have shown their capacity of generating functional AMPs from scratch with wet-lab experimental validation^14–17^. Nevertheless, these models have coupled the process of sequence representation and generation, resulting in limitations on the length, diversity or success rate of generated sequences.

Considering these existing limitations, we aim to develop a novel, user-friendly pipeline that can rapidly and efficiently generate abundant unnatural AMPs without the requirement for prior knowledge of peptide structures or amino acid sequence alignments. We develop an innovative integration of pre-trained protein language models with a class of emerging generative models, specifically diffusion models, for the design of AMPs. Pre-trained large protein language models have demonstrated their proficiency in uncovering the intricate grammar of protein sequences and extracting features from such sequences^18–23^. Diffusion models represents the zenith of generative capabilities and are excellent in the controllable generation of both images and texts^24–30^. By decoupling the tasks of sequence representation and generation, we leverage inherent strengths of both models, capturing the informative patterns by protein language model and generating novel AMPs sequences by applying diffusion models to reconstruct latent features.

Here we developed a novel and modularized deep generative model strategy, ProT-Diff, by integrating a diffusion model between the decoder and encoder of a pre-trained language model. Our ProT-Diff model proved to be effective in generating unnatural AMPs with significant diversity and a wide range of amino acid lengths. After in silico screening of thousands of generated AMPs based on physicochemical properties and predicted antimicrobial activities, we synthesized and experimentally evaluated 35 generated candidate AMPs and found 34 of them displayed efficacy against either Gram-positive or Gram-negative bacteria. In particular, we assessed one promising AMP candidate for in vivo antimicrobial activity using a lethal mouse model of acute peritonitis. Our study highlights the potential of combining pre-trained protein language models with diffusion models to generate functional peptide sequences, thereby accelerating new drug development through model-driven functional sequence design.

## Results

### Development of a deep generative model for AMPs

To address the challenges in developing AMPs by de novo design, we introduced a deep generative model which sandwiches a continuous diffusion model between the encoder and decoder of the transformer-based protein language model ProtT5-XL-UniRef50^23^. This combination allows us to benefit from ProtT5’s robust feature extraction capabilities while harnessing the diffusion model’s ability to generate continuous tensors (Fig. 1a). We decoupled the ProtT5 encoder and decoder to facilitate manipulation of peptide sequences. This decoupling allows us to project peptide sequences onto a continuous latent space using the ProtT5 encoder and then translate the generated tensors back into peptide sequences using the ProtT5 decoder. The parameters of both encoder and decoder can be frozen without further fine-tuning. To construct a generative pipeline for peptides with specific properties, such as AMPs, we only need to train a continuous diffusion model on the latent space using a well-defined peptide dataset that exhibit the desired properties. This approach maximizes the efficacy of the pre-trained protein language model and minimizes GPU memory requirements, training time, and training data needs.

**Fig. 1.**
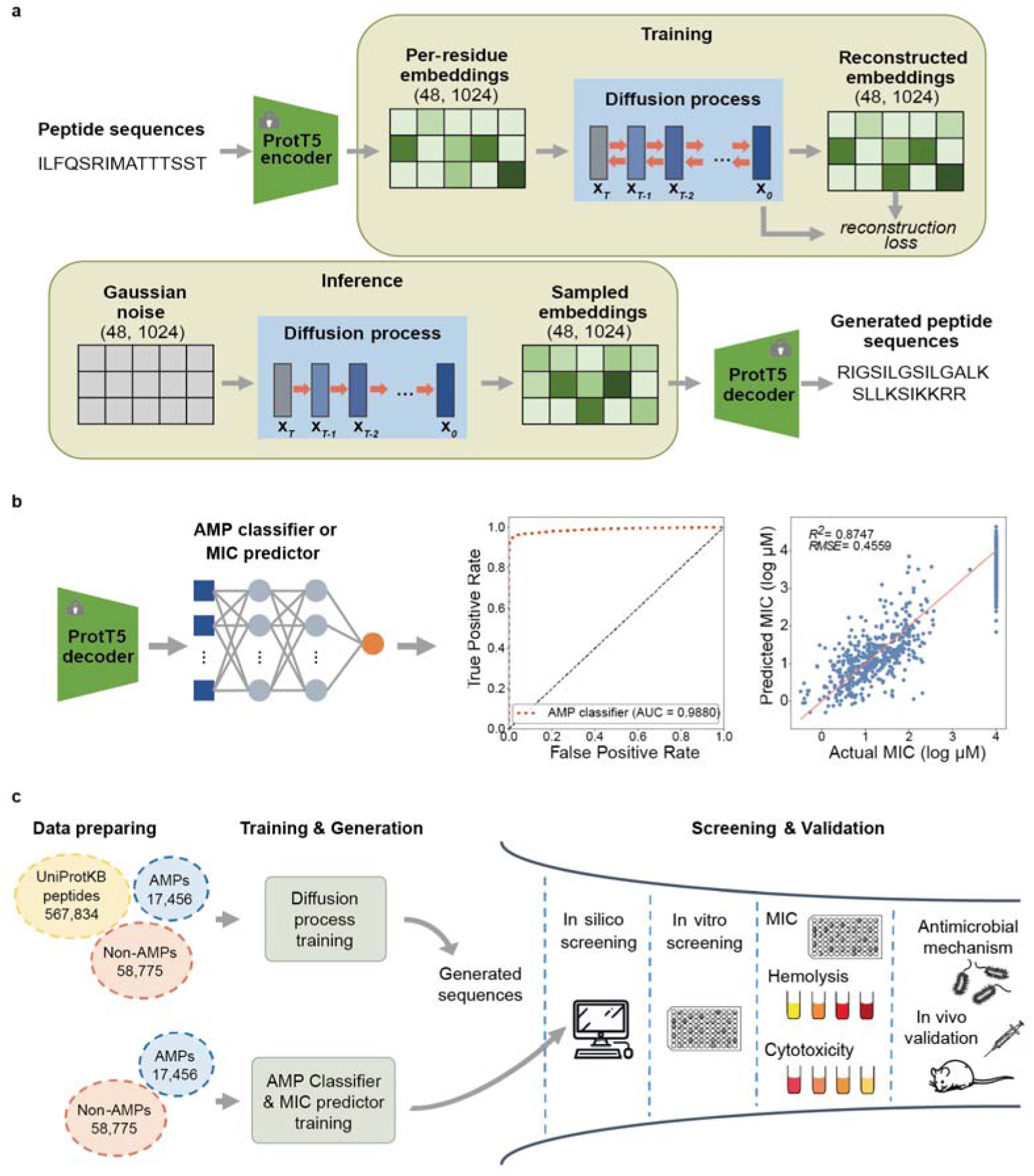
Overview of the generative pipeline for AMPs. **a**, The model architecture of ProT-Diff. ProT-Diff involves a decoupled and frozen pre-trained encoder and decoder of ProtT5-XL-UniRef50. Positioned in the middle is a diffusion model that operates on a continuous space with dimensions of (48, 1,024). This model is trained on custom peptide datasets. During training, the ProtT5 encoder embeds the input peptide sequences into the latent space, while the diffusion model is trained in a self-supervised manner to reconstruct the peptide embeddings. In the inference phase, the diffusion model denoises the initial Gaussian latent feature to generate novel peptide embeddings, which are subsequently decoded by the ProtT5 decoder into peptide sequences. **b**, The model architecture of the AMP classifier and MIC predictor, the Receiver Operating Characteristic (ROC) curve of the AMP classifier and the correlation between the actual log MIC value and the predicted log MIC value by the MIC predictor on the test set. RMSE, root mean square error. n=1,815. **c**, Workflow for generating and screening candidate AMPs. Novel peptides are generated utilizing the trained diffusion model. An AMP classifier and MIC predictor are trained in parallel. To refine the generated peptides, a series of in silico filters are applied, and the resulting candidate peptides are synthesized and subjected to both in vitro and in vivo validation.

In order to handle the variability in length of the input peptides, we padded the per-residue embeddings derived from the ProtT5 encoder to a fixed shape of (48, 1,024) with zeros. Correspondingly, the diffusion model was configured with the same fixed shape. Herein, the original peptide features generated by the diffusion model mirrored the shape (48, 1,024) of the input padded peptide embeddings. Since these generated tensors still preserve the padding patterns of the input peptide features, we removed rows that primarily consisted of values close to zero from those tensors. This truncation process enhances the decoding of these features into peptide sequences and simplifies the reconstruction of peptides with varying lengths.

During the training phase, the diffusion denoising network takes the padded peptide embeddings as input and recovers the input embeddings step by step in a self-supervised manner. In the inference phase, the latent variables are initially sampled from Gaussian distribution, and subsequently processed through the reverse steps of the trained diffusion model to obtain the denoised peptide embeddings in the ProtT5 latent space (Fig. 1a).

### Efficient generation of candidate AMPs

To generate and screen candidate AMPs, we acquired known AMPs from publicly available AMP databases. The UniProtKB peptide dataset includes substrings randomly sampled from proteins present in the UniProtKB reviewed protein database. The length distribution of this dataset matches that of known AMPs. The non-AMPs were collected from a previously published dataset^13^. We employed two strategies during the training procedure: (i) train the diffusion model with a combined dataset of known AMPs and non-AMPs; and (ii) the pretrain-finetune approach in which we initially pre-trained the diffusion model on the UniProtKB peptide dataset to learn a general grammar of protein sequences, and then fine-tuned the pre-trained model on the specific AMP dataset to capture the distinctive features of AMPs. The entire training process of the diffusion model was completed in <40 hours using a single RTX4070Ti GPU.

In parallel, we employed a three-layer multi-layer perceptron (MLP) architecture to train an AMP classifier and an AMP minimal inhibitory concentration (MIC) predictor using labeled data. These components served as filters for the generated peptides (Fig. 1b). To assess the performance of the AMP classifier, we utilized (i) the Area Under the Receiver Operating Characteristic (AUROC); and (ii) the coefficient of determination R^2^ to evaluate the goodness of fit of the AMP MIC predictor. On the test sets, the trained AMP classifier achieved an AUROC of 0.988, demonstrating an excellent performance. Furthermore, the MIC predictor yielded an R^2^ value of 0.875, indicating a high level of accuracy.

We initiated the generation process by sampling random variables from Gaussian distributions. In cases where the diffusion model was trained on a single dataset with a limited number of AMPs and non-AMPs, we introduced uniform noise distributions instead of the original Gaussian noise at each timestep to enhance the diversity of the generated content. Increasing the dispersion of the sampled noise resulted in generating more diverse tensors (Extended Data Fig. 1). In addition, in the pre-training and fine-tuning scenario, where the diffusion model was fed with ample training data, we maintained the default Gaussian noise, resulting in adequate generation diversity.

To generate enough peptides, we performed multiple iterations of the sampling procedure with different random seeds. The generated peptides were fed through a series of in silico filters. We first removed the duplicate peptides and those already present in the known AMP dataset. Subsequently, we retained only the peptides that were predicted to be AMPs by our trained classifier. Based on empirical observations, it is evident that most known AMPs (99.35%) consist of a maximum of 6 amino acids in tandem repeat. Additionally, positively charged AMPs have a higher propensity to interact with the negatively charged surfaces of bacterial membranes^31, 32^. A high positive charge is also associated with an elevated risk of hemolytic activity and cytotoxicity^7, 33^. Therefore, we imposed several constraints on the peptides, including a maximum of 6 tandem repeats of amino acids, a positive charge requirement, and a restriction of ≤40% residues being either arginine (R) or lysine (K)^33^. In subsequent sections, these generated peptides were subjected to (i) in silico filters to high-confidence candidate AMPs, (ii) in vitro validation, and (iii) in vivo validation (Fig. 1c).

### In silico assessment of candidate AMPs

We next conducted an in silico analysis to examine the physicochemical properties of the filtered candidate AMPs (Fig. 2). The analysis revealed that a majority of the generated peptides had lengths ranging from 10 to 25 amino acids, consistent with the length distribution observed in known AMPs (Fig. 2a). In addition, the filtered generated peptides displayed elevated values of net charge, isoelectric point and hydrophobic moment compared to a non-AMP dataset (Fig. 2b-e). They also demonstrated an analogous amino acid composition to known AMPs (Fig. 2g), suggesting that the trained diffusion model effectually captures the essential physicochemical properties of known AMPs that are manifested in the generated peptide set. To assess the sequence identity between known AMPs and generated AMPs, we conducted a sequence search by blastp. The sequence identities ranged from <20% to 100% (Fig. 2f), indicating that the diffusion model can not only replicate sequences present in the training set but also generate completely novel sequences. Moreover, the t-SNE projections demonstrate the model’s dual functionality in generating both known and novel sequences. The projections of the embeddings derived from the filtered generated peptides exhibited clustering with the padded peptide embeddings of known AMPs in the training set, while also exhibiting a wider distribution than the known AMPs (Fig. 2h). Collectively, these results suggest that this model can generate highly effective AMPs that do not naturally occur in biological systems.

**Fig. 2.**
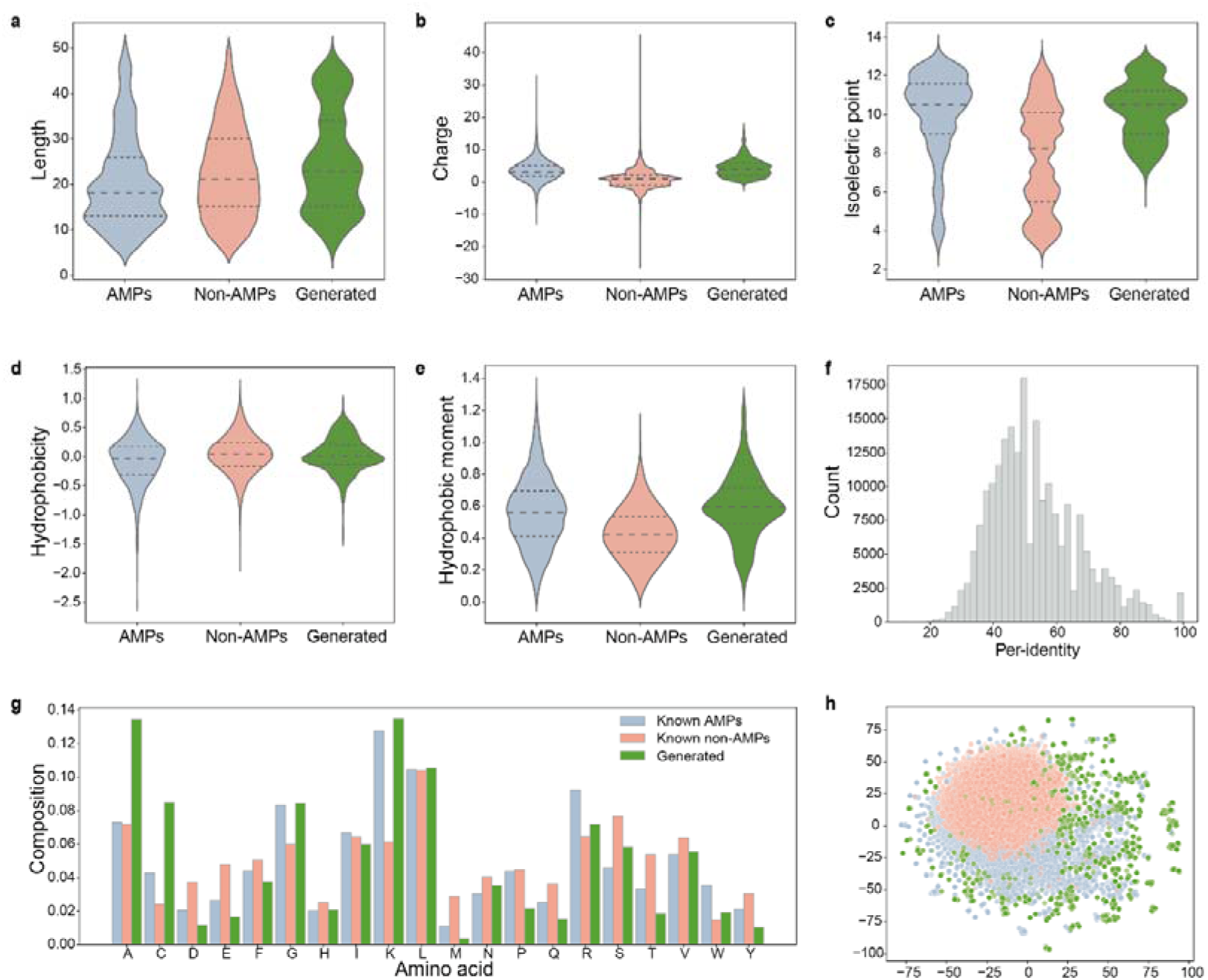
Physicochemical properties of the generated AMP candidates post in silico filtering. **a**, The amino acid lengths of peptide sequences. **b**, The theoretical net charge (pK scale = Dawson). **c**, The isoelectric point (pK scale = Dawson). **d**, The hydrophobicity index (scale = Eisenberg). **e**, The hydrophobic moment. **f**, Per-identity of the generated candidate peptides to known AMP dataset calculated by blastp. **g**, Amino acid composition. **h**, t-SNE projection of peptide embeddings. Blue dots represent known AMPs, red dots represent known non-AMPs, and green dots represent generated set. n=17,456 for AMPs, n=58,775 for non-AMPs, and n=3,133 for generated set.

### In vitro evaluation of antimicrobial activity for candidate AMPs

To evaluate the performance of our predicted AMPs, we selected 40 sequences with predicted MIC values below 10 uM for further chemical synthesis and experimental validation (Extended Data Table 1). Among them, 35 were successfully produced. We evaluated the antimicrobial activities of these 35 selected peptides against *Escherichia coli*, *Pseudomonas aeruginosa*, *Salmonella typhi* and *Staphylococcus aureus*. We assessed the antimicrobial activities by comparing the relative changes in OD_600_ between the test groups and the control group; with a threshold of 0.8 to differentiate between effective and non-effective AMPs. Strikingly, >85% (30 out of 35) of the candidate AMPs effectively inhibited at least one bacterial strain at a low concentration, and >97% (34 out of 35) of them exhibited antimicrobial activity at a high concentration (Fig. 3). Notably, most of these AMPs displayed greater antimicrobial efficacy against *E. coli* compared to other species, presumably due to the higher prevalence of AMPs targeting *E. coli* in the training set (Extended Data Fig. 2).

**Fig. 3.**
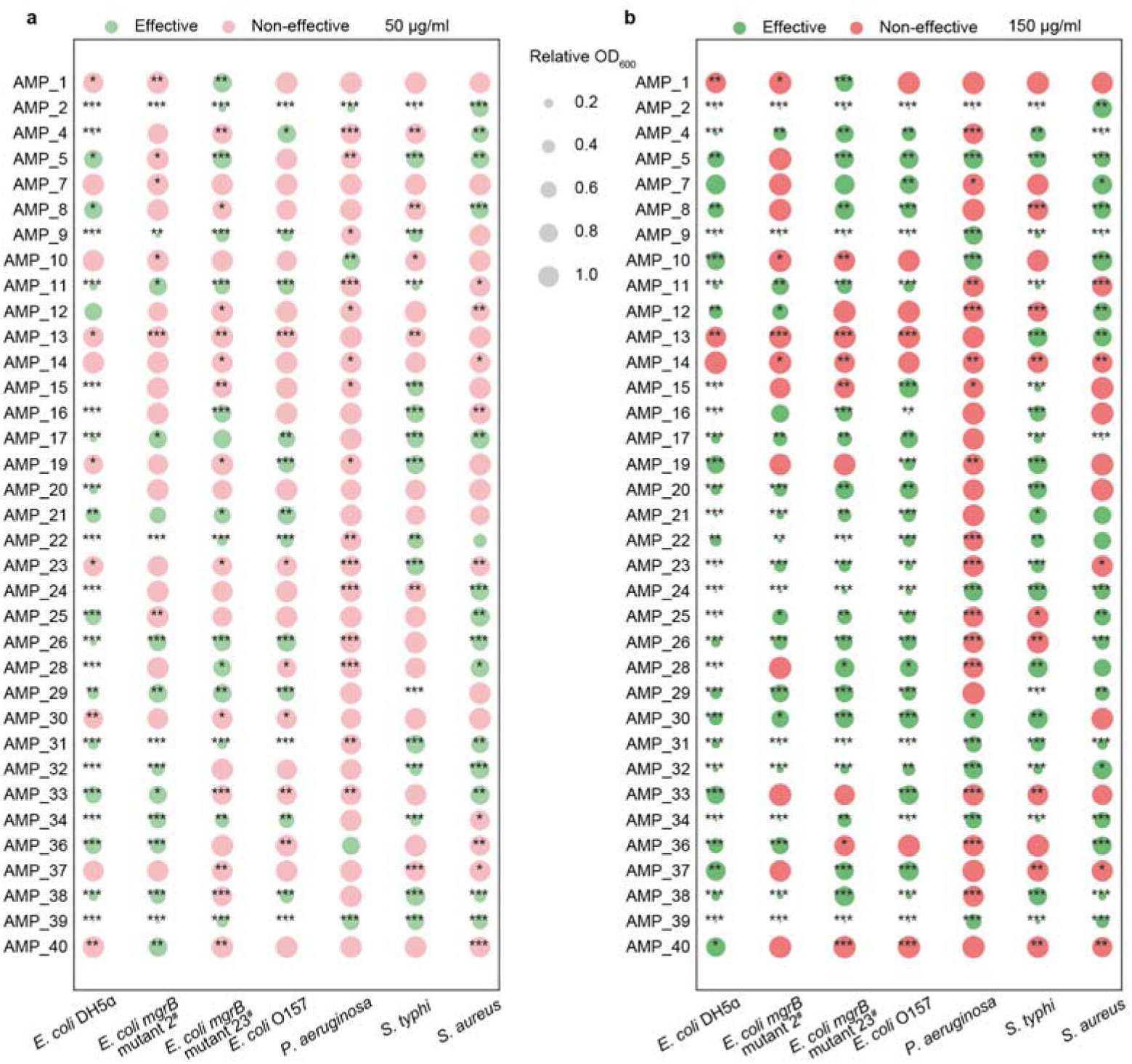
In vitro evaluation of antimicrobial activity for generated AMP candidates. The antimicrobial activities of the 35 synthesized candidate AMPs were assessed at concentrations of 50 μg/ml (**a**) and 150 μg/ml (**b**) against multiple bacteria in liquid medium. The relative OD_600_ of the experimental and control groups were compared, with an effectiveness threshold of 0.8. Any relative OD_600_ value below 0.8 was considered effective. The means between the experimental and control groups were compared using a two-sided Student’s t-test (‘*’ indicates 0.01 < *P* ≤ 0.05, ‘**’ indicates 0.001 < *P* ≤ 0.01, and ‘***’ indicates *P* ≤ 0.001). n=4 for each group.

Given our primary objective of generating novel broad-spectrum AMPs, we selected 6 synthesized AMPs (AMP_2, AMP_9, AMP_22, AMP_31, AMP_32, AMP_39) to determine their minimum inhibitory concentration (MIC), based on their excellent antimicrobial performance in the initial screening (Fig. 3). In addition to the bacteria used in the initial screening, we included two supplementary Gram-positive strains, *Bacillus sphaericus* and *Bacillus subtilis*, as well as one extra Gram-negative strain, *Acinetobacter baumannii*, in the MIC tests. The selected AMPs exhibited no discernible preference between Gram-negative and Gram-positive strains, as they demonstrated broad-spectrum activities against all the tested bacteria (Fig. 4a). All the selected AMPs consistently displayed higher MIC values (≥40 μM) when tested against *A. baumannii* and *S. aureus* compared to the other bacteria examined in this study. To verify the potential advantages of our newly developed AMP, we compared our selected AMPs with well-documented AMPs obtained from databases based on their recorded MIC values. Interestingly, no existing records document AMPs with MIC values against all the tested bacteria in our panel. Given that there are 117 known AMPs with MIC values against all the tested bacterial excluding *A. baumannii*, we compared this specific subset with our AMPs. Our AMP manifests MIC values within a moderate range, devoid of distinct favorability or unfavorability (Fig. 4b).

**Fig. 4.**
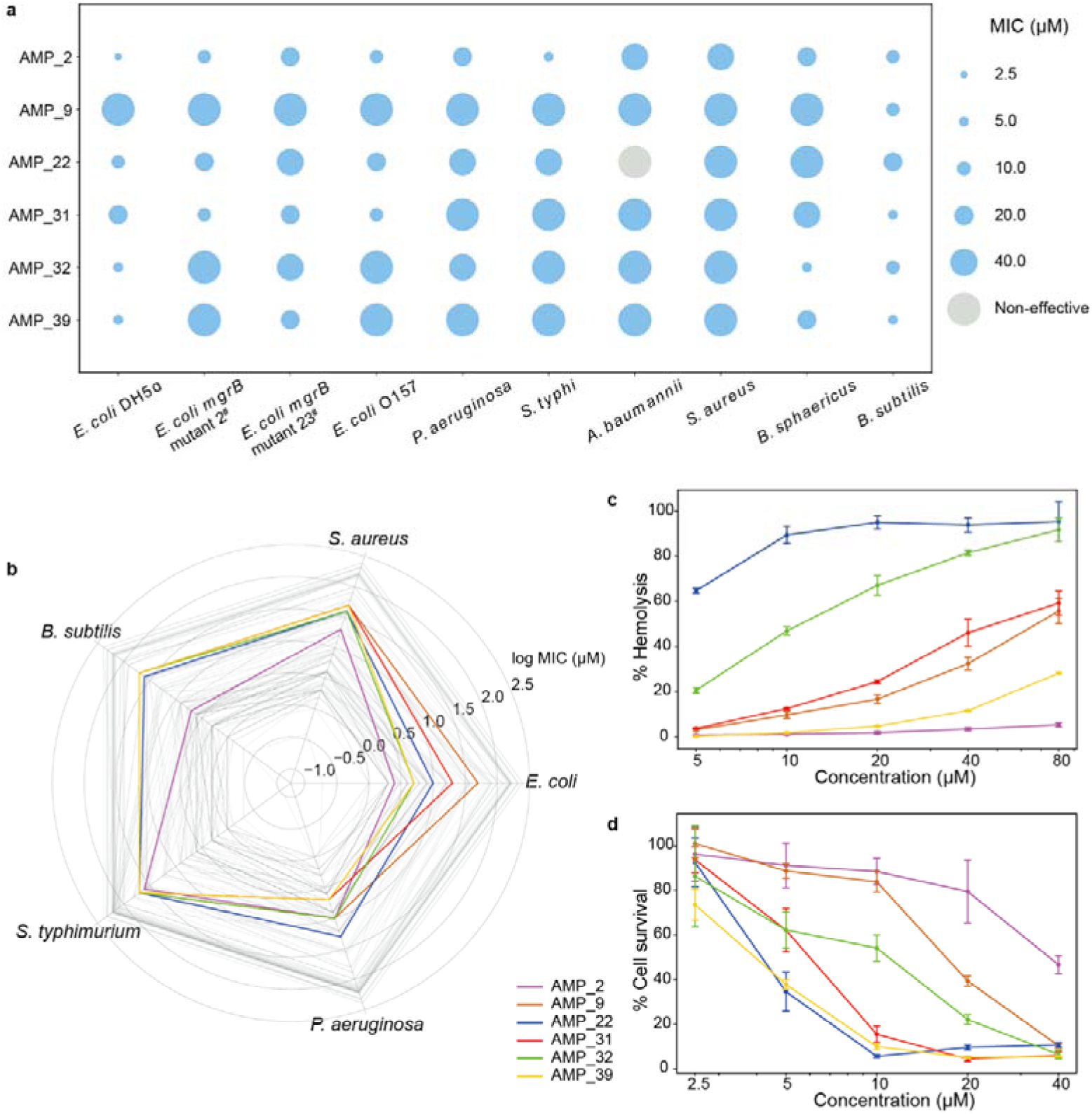
Experimental validation of generated broad-spectrum AMPs. **a**, Determination of Minimum Inhibitory Concentration (MIC) for AMPs. **b**, Comparison between our generated AMPs (colored) and well-documented AMPs obtained from databases (grey, n=117), based on their recorded MIC values. **c,d,** Assessment of hemolysis (c) and cytotoxicity (d) of AMPs. Error bars indicate standard deviation of the mean. n=3 for each group in hemolysis evaluation and n=6 for cytotoxicity evaluation.

To assess the safety of the selected candidate AMPs, we conducted evaluations of their hemolytic activity and cytotoxicity. Hemolytic activity was determined using red blood cells (RBCs) from rabbits, while cytotoxicity against human cells was assessed based on Cell Counting Kit-8 (CCK-8). Among the 6 AMP candidates, AMP_2 exhibited significantly low hemolytic activity and cytotoxicity in comparison to the other synthesized AMPs (Fig. 4c, d). Even at a concentration of 80 μM, AMP_2 did not induce noticeable hemolysis. In addition, at a concentration of 40 μM, AMP_2 maintained a cell survival rate of over 60%, signifying a half maximal inhibitory concentration (IC50) of AMP_2 higher than 40 μM. Considering the MIC values of AMP_2 against most of the tested bacteria were below 20 μM, we consider that AMP_2 showed excellent safety at its MIC value.

### In vivo evaluation of a promising, competent broad-spectrum candidate, AMP_2

As AMP_2 exhibited excellent broad-spectrum antimicrobial activity (low MIC value of 2.5 μM against *E. coli*.) coupled with low hemolytic activity and cytotoxicity, it appears to be a promising and competent broad-spectrum candidate AMP. Therefore, we next focused on this candidate for further evaluation and investigation of its antimicrobial mechanism. Since disrupting cell membrane integrity is a recognized antimicrobial mechanism, we utilized transmission electron microscopy (TEM) to observe changes in bacterial morphology after treating the bacteria with AMP_2 at its MIC for 2 to 8 hours. AMP_2 induced noticeable morphological changes in the bacteria, leading to serious leakage of cellular content (Fig. 5a). In contrast, the untreated bacterial control exhibits no apparent membrane permeabilization. Therefore, similar to certain known AMPs^34^, the action mechanism of AMP_2 involves its integration into the bacterial cell membrane, thereby disrupting membrane integrity, and culminating in cellular lysis.

**Fig. 5.**
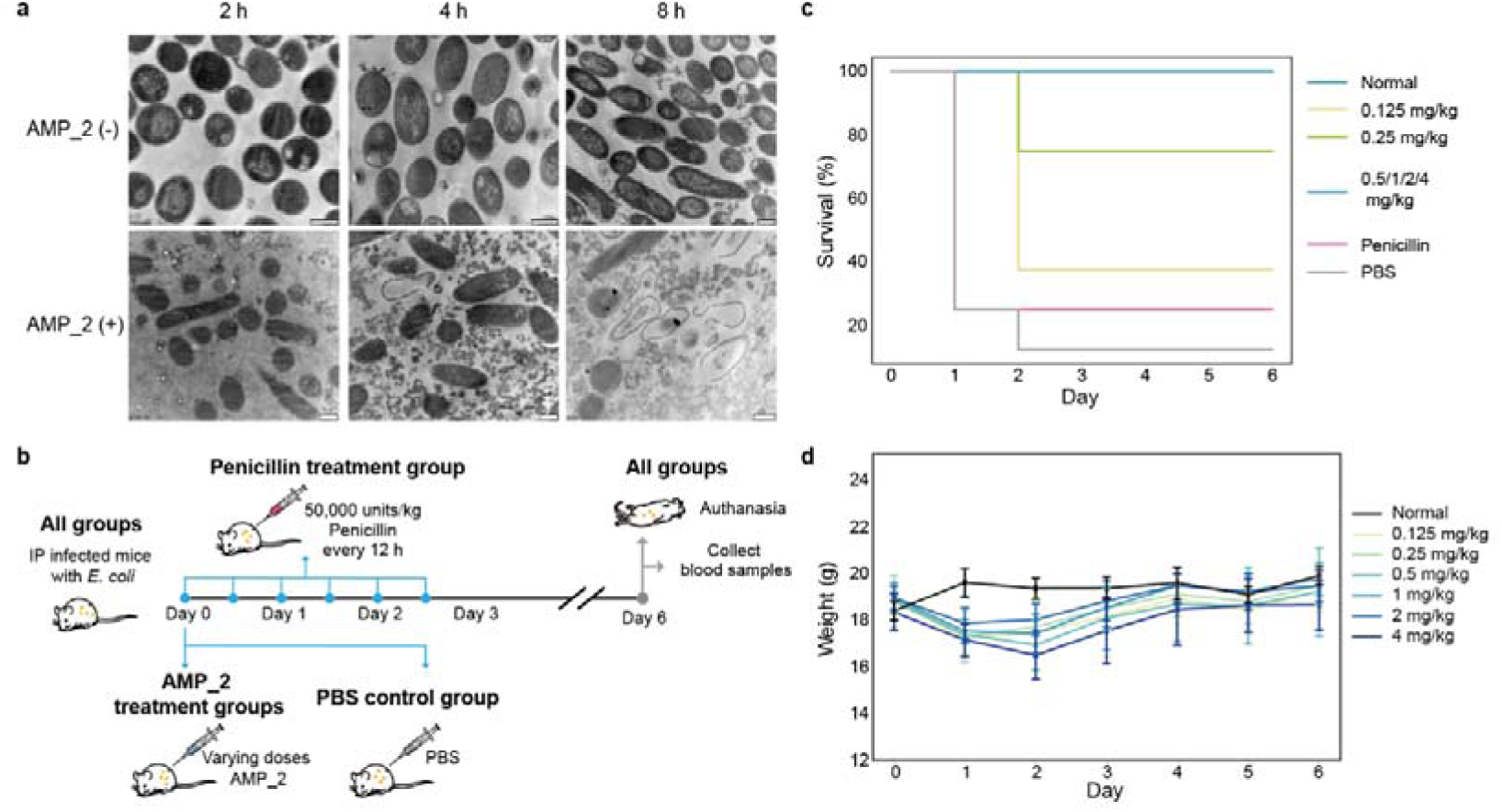
Antimicrobial mechanism and in vivo evaluation of antimicrobial activity of AMP_2. **a**, Electron microscopy images of bacteria treated without and with AMP_2 (1× MIC) at different time points. Scale bars: 500 nm. **b**, The workflow for evaluation of the efficacy of AMP_2 in a lethal mouse model of acute peritonitis. **c**, Kaplan-Meier curves showing the survival of the mice with different treatment. **d**, Monitoring the body weight of surviving mice after treatment with various concentrations of AMPs in infected mice. Error bars indicate standard deviation of the mean.

The intraperitoneal (IP) route of drug administration in laboratory animals is widely employed in numerous in vivo studies of disease model, and has been substantiated as a justifiable route for pharmacological and proof-of concept investigations^35^. Therefore, we evaluated the antimicrobial activity of AMP_2 using a well-known lethal mouse model of acute peritonitis, administered via IP route.

To establish the animal model, we employed different concentrations of the clinically isolated strain *E. coli mgrB* mutant 23^#^ to infect mice via IP injection (Fig. 5b). This strain displays multidrug resistance to a spectrum of antibiotics, including penicillin (data not shown). We found that the bacterial concentration of 5−10^5^ CFU/ml resulted in over 90% mortality within 24 hours post-infection, confirming the successful establishment of disease models in line with previous reports^36–38^. Following infection, the mice were IP administered various doses of AMP_2 immediately. Given the penicillin-resistant nature of the employed *E. coli* strain, the infected mice were treated with penicillin as a control. In accordance with the pharmaceutical company’s dosage guidance for children, a single penicillin dose of 50,000 units/kg was administered. While mice received a singular treatment of AMP_2, antibiotic treatment was administered six times at 12-hour intervals over three days. Strikingly, even at a minimal dosage of 0.5 mg/kg, AMP_2 conferred complete protection to *E. coli*-infected mice (Fig. 5c). In contrast, in the PBS treatment group, all deaths occurred within 24 hours after bacterial challenge, whereas in the other groups, deaths were observed within 48 hours. According to the log-rank test, there were significant differences between the survival of each AMP_2 treatment group and the PBS treatment group (*P* < 0.05), while there was no significant difference between the Penicillin treatment group and the PBS group (*P* > 0.05). Most surviving mice exhibited improved mental and physical well-being within 24 hours post-treatment, as evidenced by increased activity and alertness. These mice were capable of consuming food and water in a normal manner, and their fur appeared soft and glossy. Monitoring the mice’s body weight revealed a consistent increase for the surviving mice treated with AMP_2 starting from the second day post-infection (Fig. 5d). By the sixth day post-infection, while the 4 mg/kg treatment still resulted in a slight decrease in body weight, mice in the other AMP_2 treatment groups had nearly restored their body weight to that of normal mice. Furthermore, the routine blood tests revealed that many blood routine components in the treatment group did not show a significant difference compared to the normal group (Extended Data Fig. 3). Collectively, AMP_2 demonstrates promising therapeutic effects and a satisfactory safety profile in a murine model of lethal acute peritonitis.

## Discussion

In this study, we developed a deep generative approach named ProT-Diff, which ingeniously combines a protein language model and a diffusion model to generate AMPs from scratch. By decoupling the tasks of sequence representation and generation, our approach overcomes the previous limitations, especially in terms of success rate, enabling the effective and automated generation of novel AMPs. A validation of 35 selected AMP candidates revealed that 34 exhibited antimicrobial activities, highlighting the high accuracy of our approach. ProT-Diff has several advantages. First, the pre-trained protein language model ProtT5-XL-UniRef50^23^ in ProT-Diff presented strong abilities in extracting representations that reflect AMP attributes. Despite the low sequence similarity and high structural diversity among known AMPs, there was clear clustering of natural AMPs and non-AMPs in t-SNE projections of peptide embeddings (Fig. 2h). These findings, along with the high accuracy demonstrated by the AMP classifier and MIC predictor (Fig. 1b), indicating the direct production of meaningful semantic representations of protein sequences by the pre-trained language model without any fine-tuning. The excellent performance in extracting representations not only eliminates the need for prior knowledge of structures and sequence alignments, but also allows our proposed strategy to easily apply to various peptide datasets and efficiently processed on consumer-grade GPUs. Second, our diffusion model in ProT-Diff strikes a balance between diversity and fidelity in peptide sequences design, even with training sets comprising only several thousand AMPs. Diffusion models have been proven to exhibit top-notch performance in generating novel and high-quality data, particularly in data with high-dimensionality or complex structures^39^. The significant fidelity produced by diffusion model is also achieved in our study, as the length, amino acid composition and physicochemical properties, along with the clustering of the peptide embeddings in t-SNE projections present similar distributions between the generated set and the known AMPs set (Fig. 2). To overcome the challenge of maintaining diversity in generating products when the training data is limited, we optimized the noise distributions during sampling from the diffusion model or pre-train the diffusion model followed by fine-tuning. This approach is further supported by the observation that the majority of unique generated sequences are not present in the training set (Extended Data Table 2), and the lowest sequence identity between the generated sequences and training sequences is <30% (Fig. 2f). Furthermore, compared to training directly on a single peptide dataset, pre-training then fine-tuning the diffusion model leads to even greater generation diversity as well as fidelity (Extended Data Table 2).

Third, our modularized framework combines the strengths of language models and diffusion models, enhancing the generative capabilities of language models while reinforcing the representation capabilities of diffusion models. Conventional diffusion models typically operate on continuous data, whereas our task of peptide sequence generation involves discrete data. Hence, the diffusion model can operate and leverage its advantages only through the utilization of a protein language model to map discrete amino acid sequences to a high-dimensional latent space. Additionally, since the language model can naturally process variable-length sequences and the diffusion model excels in reconstructing the trained data pattern, ProT-Diff is able to generate AMPs with a wide length range. Considering the high cost and difficulty associated with solid-phase chemical synthesis of peptides of >50 amino acids, we set a maximum length of 48 amino acids in our study. Consequently, ProT-Diff enables the generation of AMPs with varying lengths up to 48 amino acids in this study. In contrast, previous AMP generation models that did not incorporate language models generated relatively short AMPs (<25 amino acids) or peptides of a fixed length^14, 15, 40^.

Among our chosen AMPs with broad-spectrum efficacy, AMP_2 demonstrated effectiveness against various drug-resistant bacteria with low MIC value and high safety. In general, a higher antimicrobial potency in antimicrobial peptides often correlates with an elevated risk of heightened cytotoxicity and hemolytic activity. For example, melittin, a well-known natural AMP, shows an exceptional antimicrobial activity, but also demonstrates high hemolysis and cytotoxicity. The MIC of melittin against *E. coli* falls within the range of 1 μM to 15 μM, comparable to our AMP_2^41–43^. However, even at a low concentration of 10 μg/ml (3.5 μM), melittin induces >50% human cell death^44^, and at 1 μg/ml (0.35 μM), it causes 50% hemolysis of cells^45^. In comparison, AMP_2 exhibited a more favorable hemolytic and cytotoxic profile, as AMP_2 induced hemolysis in <10% of RBCs at a concentration of 80 μM and resulted in approximately 50% cell death at a concentration of 40 μM (Fig. 4c, d). In particular, our in vivo experiments revealed that AMP_2 provided complete protection to mice infected with clinical drug-resistant *E.coli* strain without causing obvious damage. We observed several-fold changes in some blood routine components, such as white blood cell and lymphocyte counts, between the treatment group and the normal group (Extended Data Fig. 3) but it is commonly assumed that for murine blood components, such variations often fall within the normal physiological range. Thus, AMP_2 holds a great promise as an antimicrobial candidate deserving further investigation.

Although the toxicity of most known AMPs remains inadequately investigated, it is essential to recognize that the antibacterial efficacy of our generated AMPs using ProT-Diff does not significantly surpass that observed in natural AMPs (Fig. 4b). Up to now, the powerful deep learning methods have effectively solved the sequence-structure relationship of proteins at atomic accuracy^21, 46, 47^, enabling researchers to generate idealized protein structures^48–52^. However, de novo design of new proteins or peptides with significantly improved functionality compared to natural counterparts is still a major challenge. For the design of AMPs, there is persistent goal to create AMPs characterized by exceptionally low MIC values, minimal toxicity and other excellent performance. As the quality of the generated contents from deep generative models extensively relies on the quality of the input data, with the increasing amount and quality of training data in the future we believe this goal could be achieved through generative AI.

In summary, our study presents a lightweight and user-friendly model for efficient generation of AMPs with arbitrary length from scratch, without the necessity for prior knowledge of structure or sequence alignment. Beyond its immediate application in AMP development, this strategy is also ready to facilitate the creation of other peptide-based drug candidates in future as well as proteins with tailored characteristics. The versatility demonstrated in our approach underscores its potential impact on advancing diverse aspects of peptide and protein engineering.

## Online Methods

### Dataset

The known AMP dataset was collected from 4 public AMP databases: CAMPR4 (Collection of Anti-Microbial Peptides)^53^ (http://www.camp.bicnirrh.res.in/), ADAM (A Database of Anti-Microbial peptides)^54^ (http://bioinformatics.cs.ntou.edu.tw/adam/), APD3 (The Antimicrobial Peptide Database)^55^ (http://aps.unmc.edu), and GRAMPA (Giant Repository of AMP Activities)^33^ (https://github.com/zswitten/Antimicrobial-Peptides). The AMP records were screened based on the following criteria: 1) labeled as antibacterial, antifungal, antiviral and antimicrobial; 2) have a length of 5 to 48 amino acids; 3) only include capital letters, excluding “U, Z, O, B, J” residues. A total of 17,456 known AMPs were collected by combining these 4 databases and removing duplicate sequences.

The truncated UniProtKB Reviewed protein dataset was created by truncating sequences from the UniProtKB Reviewed protein database based on the length distribution of the known AMP dataset. A total of 567,834 peptide sequences were collected after removing duplicates. The non-AMPs containing 58,775 peptides were collected from a previously published dataset^13^. The training and test sets of various training tasks were consistently divided in a ratio of 8:2.

### The overview of the ProT-Diff generation model

We first embedded the peptide sequences in the training and test sets into tensors of fixed shape by a pre-trained language model for proteins. The peptide embedding features were utilized to independently train a diffusion process on continuous space, which enabled us to sample from the trained diffusion process to get novel peptide features in the embedding space. Subsequently, the decoder of the pre-trained language model was employed to decode the generated peptide features and retrieve amino acid sequences. From the generated peptide sequences, only those predicted as AMPs by the AMP classifier and adhering to the predetermined physicochemical property constraints were considered as candidate novel AMPs.

### Peptide sequence embedding

We selected the ProtT5-XL-UniRef50 encoder of the pre-trained protein language models from ProtTrans to generate embeddings for peptide sequences^23^. The resulting residue embeddings have a shape of (peptide length, 1,024) and are padded with zeros to a shape of (48, 1,024) before being inputted into the subsequent diffusion process.

### Training the diffusion process

The training procedure (Extended Data Fig. 4a) for the diffusion process follows the framework of Diffusion-LM^29^. The diffusion model was trained directly on the known AMPs combined with non-AMPs, or by the pretrain-finetune approach. For the latter scenario, we first pre-trained the diffusion process on the truncated UniProtKB Reviewed protein dataset containing 567,834 peptides combined with 17,456 known AMPs, and then fine-tuned the diffusion process on the known AMPs dataset. The sample weights of UniProtKB peptides and the known AMPs during pre-training were set to 0.52 and 16.78, respectively, according to the size of the datasets.

The number of diffusion steps during training was set to 2,000. The Trans-UNet architecture was employed to predict **x_O_**, and the noise schedule was set to *sqrt (square-root)* schedule proposed by Diffusion-LM^29^. The total number of parameters of the diffusion model was 20,706,816.

### Sampling for peptide features

DDPM sampler was applied to sample peptide features in the embedding space. The diffusion steps of generative diffusion process were down sampled from 2,000 steps to 200 steps according to Diffusion-LM^29^. The noise of each diffusion step was sampled from either a normal distribution by default or uniform distributions. The sampling algorithm is present in Extended Data Fig. 4b.

### Decoding the generated features

The generated peptide residue embeddings were wrapped as encoder outputs and were passed to the ProtT5-XL-UniRef50 decoder to reconstruct the amino acid sequences.

### AMP classifier

The AMP classifier, used to distinguish AMPs from non-AMPs, is implemented as a simple multilayer perceptron (MLP) with 3 fully-connected layers. The classifier model reads the peptide residue embeddings of ProtT5-XL-UniRef50 padded to shape (48, 1,024) as inputs and outputs the classification results. The classifier was trained using the residue embeddings of known AMPs and non-AMPs with binary labels. The positive training set of known AMPs was collected from CAMP, ADAM and APD3 databases, and the negative training set of non-AMPs was obtained from the previous study c_AMP-prediction^13^. The dropout rate of each hidden layer was set to 0.2 and the L2 regularization factor was set to 0.001.

### AMP MIC predictor

The architecture of the AMP MIC predictor is identical to that of the AMP classifier, with the exception of the activation function used in the output layer. The MIC predictor is a regression model trained with log MIC values of known AMPs from GRAMPA database^33^. For AMPs with multiple targets measurements, we first took the geometric mean of all the MIC values and then took log to get the final log MIC value.

### Pre-screening for candidate novel AMPs

The generated set of AMP sequences was sieved with the following filters: (i) removing duplicates; (ii) eliminating sequences already present in the known AMPs set; (iii) discarding the sequences predicted as non-AMPs by the AMP classifier; (iv) excluding sequences containing more than 6 tandem repeat amino acids; (v) removing the sequences with a non-positive charge; and (vi) excluding sequences with a proportion of lysine (K) and arginine (R) residues exceeding 40%. The remaining generated AMPs that met all these constraints could be regarded as candidate novel AMPs and be passed to further experimental validations.

Of the 40 selected candidate peptides that were synthesized, AMP_1–20 were produced by the first training approach, while AMP_21–40 were produced by the second training approach.

### Peptide synthesis

Peptides were prepared by solid-phase synthesis with purity of >90% by Genscript Biotech Corporation (Nanjing, China). The net peptide content was quantified by analysis of nitrogen content. Peptides as Trifluoroacetic acid (TFA) salts were used in antimicrobial activity test. TFA in peptides were removed by replacement with acetate for the determination of minimal inhibition concentration (MIC) and animal tests.

### Bacterial strains

Gram-negative bacteria include *E. coli* DH5α, *E. coli* O157: H7 CICC 21530, *S. typhimurium* CICC 21484, *A. baumannii* AB6 and *P. aeruginosa* ATCC 15442. Gram-positive bacteria include *S. aureus* ATCC 33591, MRSA ATCC 43300, *B. subtilis* ATCC 9372. *L. sphaericus* CCM 2177. Two *E. coli* clinical isolates with *mgrB* mutation from Tongji Hospital (Wuhan, China) were used in this study, namely *E. coli mgrB* mutant 2^#^ and *E. coli mgrB* mutant 23^#^.

### Antimicrobial activity test

The antimicrobial activity test was performed as described previously^56^. In brief, the procedure involved the following steps.

Firstly, the tested bacteria were streaked on Luriae–Bertani (LB) agar medium and incubated at 37°C overnight. Individual colonies were selected from the agar plate and transferred to Mueller–Hinton broth (MHB) for further cultivation. The culture was shaken at 160 rpm at 37°C overnight. Subsequently, the bacterial suspension was transferred to fresh MHB at a ratio of 1:100 and incubated at 37°C. When the optical density at 600 nm (OD_600_) of bacterial suspension reached 0.6–0.8, the bacterial suspension was further diluted with MHB to OD_600_ of approximately 0.1.

For the preparation of the antimicrobial peptide solution, the AMP was dissolved in either sterile water or dimethyl sulfoxide (DMSO) to achieve a concentration of 10 mg/ml. This solution was further diluted with MHB medium to attain the desired concentration for testing.

The antibacterial activity tests were conducted in 96-well plates, and five experimental groups were established as follows : 1) blank control group, MHB solution; 2) without AMPs group, 100 μl of bacterial solution and 100 ul of MHB; 2) AMPs experiment group 1 (low concentration of AMPs, 50 μg/ml), 100 μl of bacterial solution and 100 ul of 100 μg/ml AMP; 3) AMPs experiment group 2 (high concentration of AMPs, 150 μg/ml), 100 μl of bacterial solution and 100 ul of 300 μg/ml AMP; 4) low concentration of AMPs control, 100 μl of MHB solution and 100 ul of 100 μg/ml AMP; 5) high concentration of AMPs control, 100 μl of MHB solution and 100 ul of 300 μg/ml AMP. Before incubation and after incubation at 37°C for 16 h, the OD_600_ value of each well was measured, namely OD_600 (0 h)_ and OD_600 (16 h)_. The relative OD was calculated as AMPs experiment group △OD_600_/ without AMPs group △OD_600_.

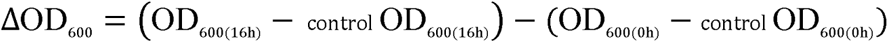

For the MIC determination of selected AMPs, 100 ul bacterial culture with an OD_600_ value of approximately 0.1 was incubated with AMPs at concentrations ranging from 0 µM to 100 µM at 37°C for 15–18 h. Herein, the MIC was determined as the minimum concentration of AMPs at which no detectable bacterial growth was observed.

All experiments were conducted with a minimum of four independent replicates. The Student’s t test was used to compare the means between experimental groups and control groups (two-sided).

### TEM measurement

Transmission Electron Microscopy (TEM) was employed to assess the cell membrane damage induced by antimicrobial peptides (AMPs). Firstly, 50 μl of *E. coli* DH5α culture with an OD_600_ of approximately 0.1 were mixed with 50 μl of AMPs solution (at 1× MIC) in a 96-well plate. This mixture was incubated at 37°C for 2 h, 4 h, 8 h and 16 h. Following the incubation period, the bacteria were pelleted by centrifugation at 6,000 rpm for 5 min and fixed with 2.5% (vol/vol) glutaraldehyde in phosphate buffer (PB, 0.1 M, pH 7.4). Subsequently, the cells were postfixed with 1% (wt/vol) osmium tetraoxide in PB for 2 h at 4°C, and were dehydrated through a graded ethanol series (30%, 50%, 70%, 80%, 90%, 100%, with each step lasting 7 minutes) before being put in pure acetone by two 10-minute steps. Following dehydration, the samples were subjected to infiltration using progressively mixed combinations of acetone and SPI-PON812 resin (composed of 16.2 g SPI-PON812, 10 g DDSA, and 8.9 g NMA) in ratios of 3:1, 1:1, and 1:3. The infiltration medium was replaced with pure resin. Finally, the cells were embedded in pure resin containing 1.5% BDMA and polymerized for 12 h at 45°C, followed by an additional 48 h polymerization at 60°C. The ultrathin sections (70 nm thick) were obtained using a microtome (Leica EM UC6), double-stained with uranyl acetate and lead citrate, and examined using a transmission electron microscope (FEI Tecnai Spirit 120 kV).

### Hemolytic activity test

The hemolytic activity of AMPs was assessed using red blood cells (RBCs) from rabbits. Initially, fresh RBCs were washed three times with phosphate-buffered saline (PBS) through centrifugation for 15 min at 1,000 g until the supernatant became clear. Subsequently, RBCs were resuspended in PBS to attain a final erythrocyte concentration of 4% (v/v). Next, 100 μl suspension of RBCs was incubated with 50 μl of AMPs solution at various concentrations at 37°C for 1 h in 96-cell cell plates. Following incubation, the supernatant was collected by centrifugation for 15 min at 1000 g and its absorbance was measured at 540 nm. For reference, the hemolysis of RBCs in PBS was designated as representing zero hemolysis, while the hemolysis of RBCs in 0.1% (w/v) Triton X-100 was considered as 100% hemolysis. The percentage of hemolysis was calculated using the following Equation.

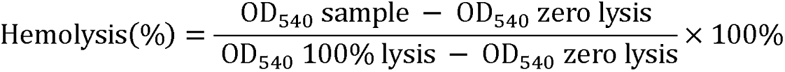

### Cytotoxicity against mammalian cells

Cell Counting Kit 8 (CCK-8) assay was employed to assess the cytotoxicity of AMPs against mammalian cells. Specifically, approximately 4,000 293T cells were seeded per well in a 96-well cell culture plate (Corning, US), and were incubated at 37°C with 5% CO_2_ for 24 h. Subsequently, the culture medium was replaced by different concentrations of AMPs solution diluted with the medium. The cells were co-incubated with AMPs solution for an additional 24 h at 37°C. And then, 10 μl of the CCK-8 reagent (Beijing Lablead Biotech, China) was added to each well for 2 h of incubation at 37°C. Finally, OD at 450 nm was measured using an imaging multimode microplate reader (Cytation 3, BioTek, US). Cell viability was defined as the percentage of each concentration accounted for of the control. Six replicates were made for CCK-8 assay.

### Murine acute peritonitis model

Female Balb/c mice, approximately 18 g in weight and aged 5–6 weeks, were used to build lethal murine acute peritonitis model according to previously established protocols^36–38^. The mice were subjected to intraperitoneal injections with *E. coli mgrB* mutant 23^#^ suspension (0.2 ml per mouse) at concentration of 5×10^5^ CFU/ml, 5×10^6^ CFU/ml, 5×10^7^ CFU/ml, respectively. After infection, the mice were observed for 24 h to determine the mortality rate. Each group consisted of 8 mice.

To access the impact of AMP_2 on acute peritonitis, the female Balb/c mice were firstly inoculated with an *E. coli mgrB* mutant 23^#^ suspension at concentration of 5×10^5^ CFU/ml, which resulted in a mortality rate of 90–100% within 24 h. Subsequently, the mice were administered varying doses of AMP_2 (0.125 mg/kg, 0.25 mg/kg, 0.5 mg/kg, 1 mg/kg, 2 mg/kg, 4 mg/kg) immediately after infection. As controls, the infected mice received intraperitoneal injections of sterile PBS or underwent penicillin treatment (0.2 ml per mouse). The single penicillin dose administered was 50,000 units/kg, and the mice received antibiotic treatment every 12 hours for three days. The behavior and survival of the animals were monitored over a 7-day period. Blood samples were collected from the mice prior to euthanasia for routine blood tests. The experiments were conducted in groups, each comprising 8 mice.

All animal experiments were conducted in ABSL-2. The animal use protocol listed in this study were approved by the Animal Experimentation Ethics Committee of National Vaccine & Serum Institute (NVSI) of Sinopharm (NVSI-RCD-JSDW-SER-2023025), according to China’s Guidelines on Welfare and Ethical Review for Laboratory Animals.

## Code availability

The ProT-Diff codes are available upon request.

## Acknowledgements

We extend our sincere gratitude to Can Peng from the Center for Biological Imaging (CBI), IBP-CAS, for invaluable technical support with the transmission electron microscopy (TEM) work. We also express our appreciation to Chao-Xiang An from the National Vaccine & Serum Institute (NVSI) of Sinopharm for his assistance with the animal experiments. Additionally, we would like to thank Jiao-Yu Deng from the Wuhan Institute of Virology, CAS, for generously providing clinically isolated strains and offering valuable insights into minimum inhibitory concentration (MIC) measurement techniques. This work was supported by the National Key Research and Development Program of China (Grant No.2022YFC2303501, No.2018YFA0902702), and by National Natural Science Foundation of China (Grant No.32271489).

## Notes

### Competing Interest Statement

The authors have declared no competing interest.

