## Extended Data Fig. 1, Extended Data Fig. 2, Extended Data Fig. 3, Extended Data Fig. 4 for "ProT-Diff: A Modularized and Efficient Approach to De Novo Generation of Antimicrobial Peptide Sequences through Integration of Protein Language Model and Diffusion Model"

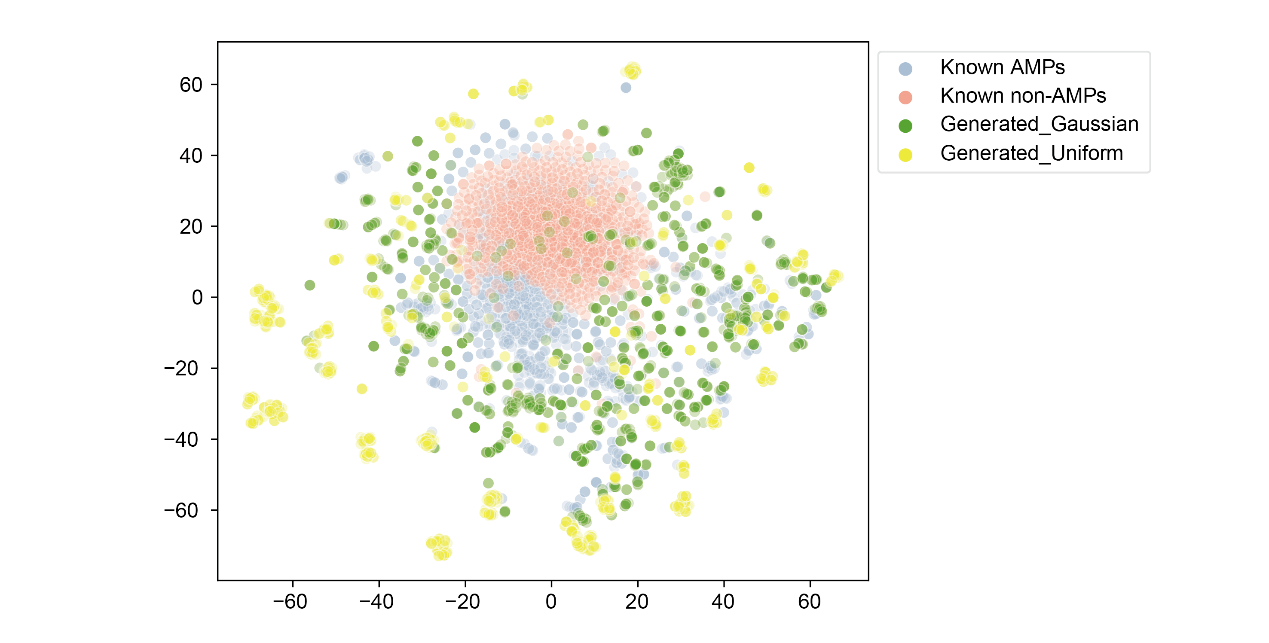

**Extended Data Fig. 1 | Comparison of generation diversity using different noise distribution during sampling from the diffusion model.** The t-SNE projection displays the peptide embeddings of known AMPs (blue) and non-AMPs (red) in the training set, as well as the generated peptides resulting from the sampling process with Gaussian noise distribution (green), and Uniform noise distribution (yellow). n=1,000 for each group.

Extended Data Fig. 2

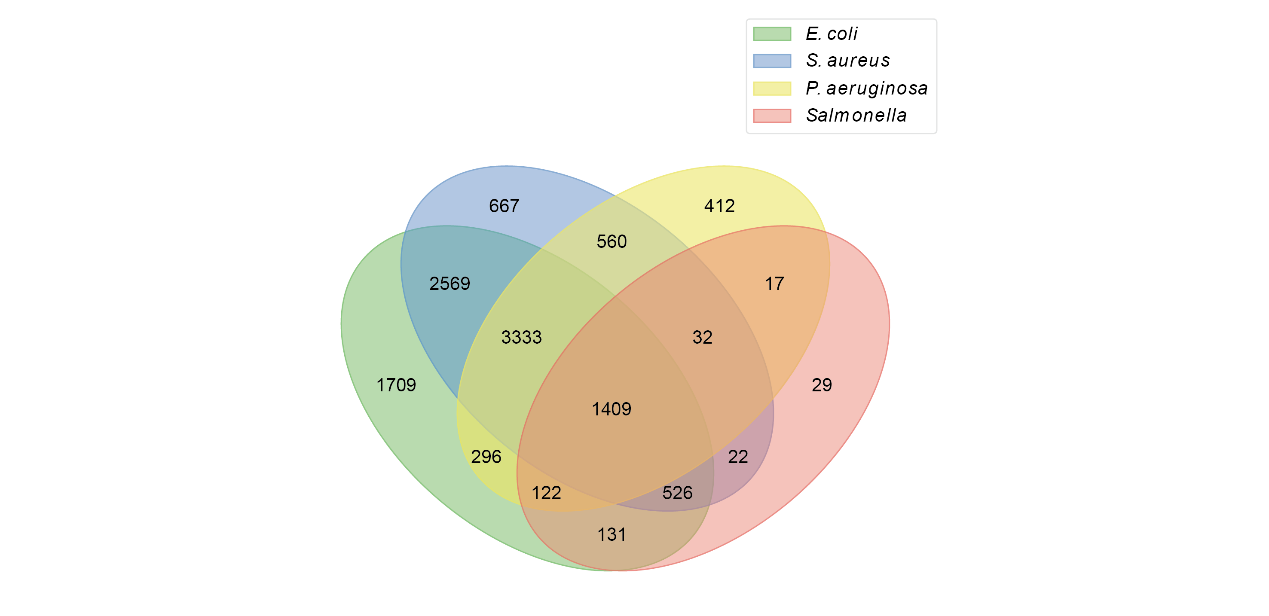

**Extended Data Fig. 2 | The presence of species target labels in known AMPs.** The overlap of known AMPs targeting *E. coli*, *S. aureus*, *P. aeruginosa*, and *Salmonella*.

Extended Data Fig. 3

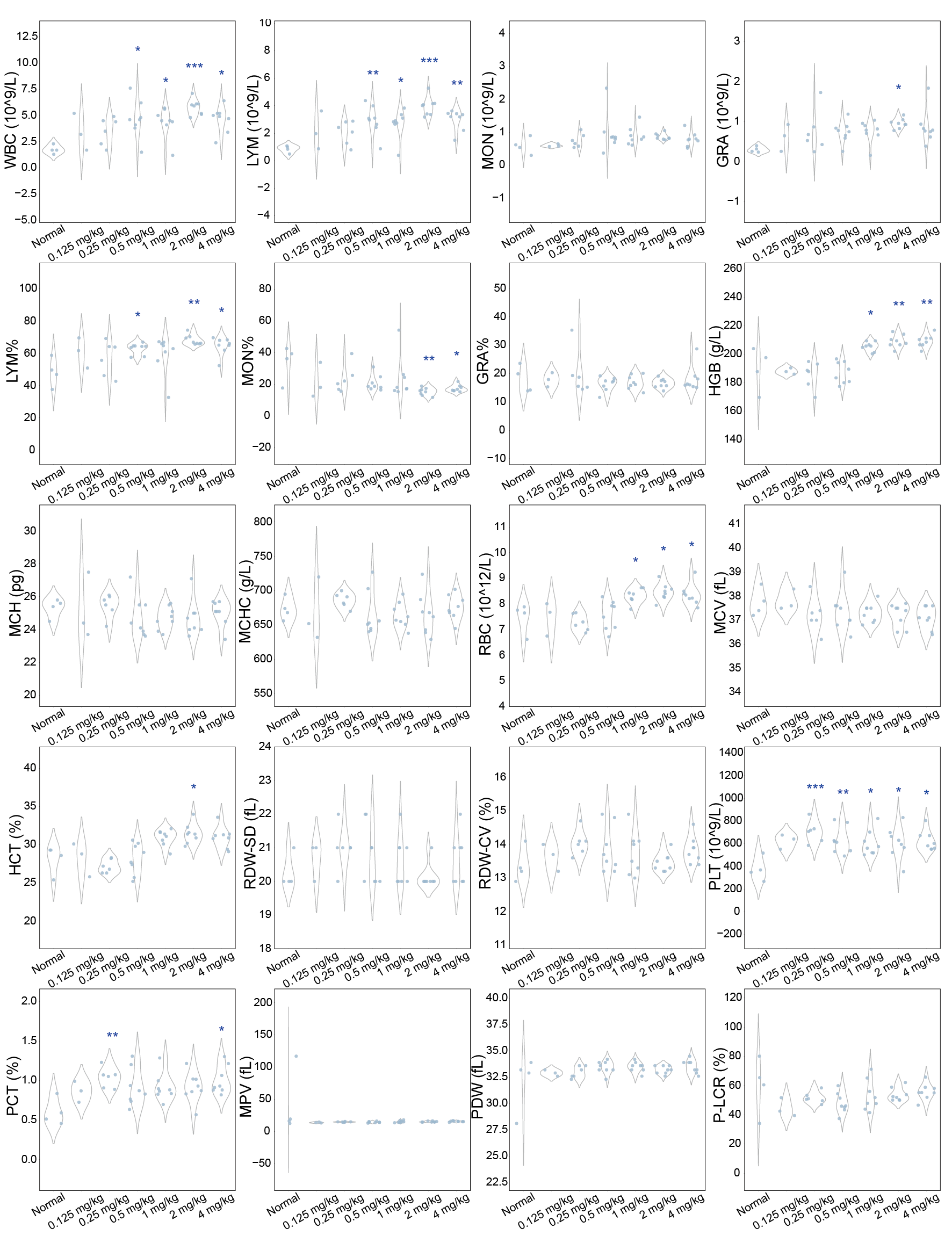

**Extended Data Fig. 3 | Routine blood analysis of mice following treatment with AMP_2 during infection.** WBC, white blood cells; LYM, lymphocytes; MON, monocytes; GRA, granulocytes; HGB, hemoglobin; MCH, mean corpuscular hemoglobin; MCHC, mean corpuscular hemoglobin concentration; RBC, red blood cells; MCV, mean corpuscular volume; HCT, hematocrit; RDW-CV, red cell distribution width reported statistically as coefficient of variation; RDW-SD, RDW reported statistically as coefficient of standard deviation; PLT, platelet; PCT, procalcitonin; MPV, mean platelet volume; PDW, platelet distribution width; P-LCR, platelet-large cell ratio. The means between the experimental and normal groups were compared using a two-sided Student’s t-test (‘*’ indicates 0.01 < *P* ≤ 0.05, ‘**’ indicates 0.001 < *P* ≤ 0.01, and ‘***’ indicates *P* ≤ 0.001).

Extended Data Fig. 4

| **a** Diffusion Training | |
| --- | --- |
| Require: Peptide sequences $\mathbf{w}$, AMP binary label $y$, Peptide embeddings $\mathcal{D}$,  Original data sample $\mathbf{x}_{\mathbf{0}}$, Latent variables $\mathbf{x}_{\boldsymbol{1}}\boldsymbol{,\ldots,}\mathbf{x}_{\boldsymbol{T}}$,  Protein sequence encoder $E\text{()}$, Denoising model $\boldsymbol{\mu}_{\boldsymbol{\theta}}\text{()}$,  Timesteps $T$, Noise $\boldsymbol{\epsilon}$, Noise schedule $\bar{\alpha_{t}}$, Learning rate $\text{η}$ | |
| 1: **repeat** |  |
| 2: $\mathbf{x}_{\mathbf{0}}\boldsymbol{=}E\text{(}\mathbf{w}\text{)}\boldsymbol{\sim}\mathcal{D}$ | Encode sampled data |
| 3: $t \sim\mathcal{U}\text{(}\{1,\ldots, T\}\text{)}$ | Sample timestep |
| 4: $\boldsymbol{\epsilon}\mathcal{\sim N}\text{(}0,\mathbf{I}\text{)}$ | Sample noise |
| 5: $\mathbf{x}_{\boldsymbol{t}}\boldsymbol{=}\sqrt{\bar{\alpha}_{t}}\mathbf{x}_{\mathbf{0}}\boldsymbol{+}\sqrt{1\boldsymbol{-}\bar{\alpha}_{t}}\boldsymbol{\epsilon}$ | Compute latent variables |
| 6: $\mathcal{L=}\left\Vert\boldsymbol{\mu}_{\boldsymbol{\theta}}\text{(}\mathbf{x}_{\boldsymbol{t}},t\text{)}-\mathbf{x}_{\boldsymbol{0}} \right\Vert_{\text{2}}^{2}$ | Compute loss |
| 7: $\theta\text{=}\theta-\text{η}\nabla_{\theta}\mathcal{L}$ | Update parameters |
| 8: **until** converged |  |

| **b** Diffusion Sampling | |
| --- | --- |
| Require: Estimated original data ${\tilde{\mathbf{x}}}_{\mathbf{0}}$, Latent variables $\mathbf{x}_{\text{1}}\boldsymbol{,\ldots,}\mathbf{x}_{\boldsymbol{T}}$,  Denoising model $\boldsymbol{\mu}_{\boldsymbol{\theta}}\text{()}$, Timesteps $T$, Noise $\boldsymbol{\epsilon}$, Noise schedule $\bar{\alpha_{t}}$,  Protein sequence decoder$\text{ }D\text{()}$ | |
| 1: $\mathbf{x}_{\boldsymbol{T}}\mathcal{\sim N}\text{(}0,\mathbf{I}\text{)}$ | Sample the initial latent variable |
| 2: **for** $t=T,\ldots,1$ **do** |  |
| 3: ${\tilde{\mathbf{x}}}_{\mathbf{0}}\boldsymbol{=}\boldsymbol{\mu}_{\boldsymbol{\theta}}\text{(}\mathbf{x}_{\boldsymbol{t}},t\text{)}$ | Estimate original data $\mathbf{x}_{\mathbf{0}}$ |
| 4: **if** $t=1$ **then** |  |
| 5: **return** ${\tilde{\mathbf{x}}}_{\mathbf{0}}$ | Get generated data |
| 6: **end if** |  |
| 7: $\boldsymbol{\epsilon}\mathcal{\sim N}\text{(}0, \mathbf{I}\text{)}\mathrm{or}\mathcal{U}\text{(}-3, 3\text{)}$ | Sample noise |
| 8: $\mathbf{x}_{\boldsymbol{t-1}}\boldsymbol{=}\sqrt{\bar{\alpha}_{t-1}}{\tilde{\mathbf{x}}}_{\mathbf{0}}\boldsymbol{+}\sqrt{1\boldsymbol{-}\bar{\alpha}_{t-1}}\boldsymbol{\epsilon}$ | Compute the next latent variable |
| 9: **end for** |  |
| 10: $\text{w}\boldsymbol{'=}D\text{(}{\tilde{\mathbf{x}}}_{\mathbf{0}}\text{)}$ | Decode generated data |

**Extended Data Fig. 4 | Diffusion training (a) and sampling (b) algorithms.**
